# Effects of short-term isolation on social animals’ behavior: an experimental case study of Japanese macaque

**DOI:** 10.1101/2021.03.28.437096

**Authors:** T Morita, A Toyoda, S Aisu, A Kaneko, N Suda-Hashimoto, I Adachi, I Matsuda, H Koda

## Abstract

One of the goals in animal socioecology is to understand the functions and dynamics of group living. While observations of free-ranging animals are a crucial source of information, an experimental investigation that manipulates the size or composition, or both, of animal groups in captivity can also bring complementary contributions to the research inquiry. When paired with an automatic data collection by biologging technology, experimental studies on captive animals also allow for big data analyses based on recent machine learning techniques. As an initial exploration of this research paradigm, the present study inquired to what extent isolation of captive Japanese macaques (*Macaca fuscata*) changed their movement patterns. Using three-dimensional location trajectories of the macaques that were systematically collected via Bluetooth Low Energy beacons and a deep neural network, we estimated the identifiability of whether a macaque was behaving in isolation or in group. We found that the neural network identified the isolation vs. in-group conditions with more than 90% accuracy from a five-minute location trajectory, suggesting that the isolation caused notable changes from the canonical group-living behaviors. In addition, the isolation made each individual more identifiable from one another based on their location trajectories.

## Introduction

Many animals live in bonded social groups that differ in size, composition, and cohesion [1,2]. To better understand proximate mechanisms behind social systems of animals as well as evolutionary factors deriving the systems, researchers have studied the costs or benefits, or boths, of group living [3–5]. For example, living in a large group may reduce predation risk while competition among members for food resources would become more intensive, as observed in various animal taxa, including primates [6], carnivores [7], birds [8], and fishes [9]. One way of addressing this research question is to observe free-ranging animals that regularly split into smaller groups (subgroups) during the day (called the fission-fusion social system). As an example, the correlation between the size of subgroups and different behavioral strategies taken by their members has been studied on many mammals, including hyenas [10,11], dolphins [12,13], and primates [14–16]. However, collecting data from free-ranging animals is challenging due to many uncontrollable environmental factors. Specifically, there is little hope to exhaustively observe all subgroups that are theoretically possible or of interest, as some of them may not be formed spontaneously. Also, observations of free-ranging animals are susceptible to the inevitable biases of the observers. In contrast, studies of captive animals make the size and composition of subgroups more controllable via experimental separation/isolation of group members. Thus, experiments in a captive environment can provide complementary contributions and a deeper understanding of group living. Similar experiments are common in animal psychology and brain science; for example, social isolation has been used to study the effects of early rearing conditions on subsequent social behaviors, specifically in depth for rodents and primates [17].

Although it is challenging to target specific group composition or sizes in free-ranging animals, quantitative difficulty in collecting their behavioral records has largely been resolved due to recent developments in biologging technology. For an instance, we can track movements of individual free-ranging animals using high-resolution global positioning satelites (GPS). The collected data can be used to study patterns in group movement, decision making, and ecological mechanisms behind them [18–23]. Biologging data are systematic, fine-grained, and large-scale as compared to traditional ecological observations. Therefore, they are ideal for machine learning-based [24–26], graph-theoretic [27–29], and other data-scientific analyses [30,31]. While biologging technology has mainly been used to study spontaneous behaviors of animals in non-controlled settings, it can also be a powerful tool to collect large-scale animal data under experimental conditions, the effectiveness of which was explored in this study.

As an initial exploration of the inquiry into the effects of social modifications (e.g, isolation/separation) on animal behavioral patterns, this study performed a quantitative evaluation of changes in movement patterns of captive Japanese macaques (*Macaca fuscata*) experimentally caused by their visual isolation. Specifically, we trained an artificial neural network to estimate the identifiability of whether a macaque was behaving in isolation or in group based on their location trajectories. The high identifiability of isolation would indicate that the macaques exhibited some movement patterns characterizing the isolation, and using machine learning techniques, such characteristic movements can also be visualized [32]. The location records of each macaque were automatically collected in three dimensions (3D) using a recently developed biologging technology. Besides measuring the extent of overall behavioral changes upon isolation, we also investigated whether isolation intensifies or diminishes the macaques’ individuality reflected in their movement, performing individual recognition based on the location trajectories [33]. On the one hand, if group-living restricts behavioral patterns, it is possible that isolation increases the individual recognition score by removing the bounds. For example, Japanese macaques form matrilineal groups based on social bonding among females, and these are known to exhibit greater spatial cohesion than males in general [33,34]. Thus, location trajectories of female macaques can look similar to one another, which would make the movement-based individual recognition difficult. However, it is also possible that isolation decreases the individual recognition score if each individual plays a specific role in the group, which reflects their individuality. In such cases, isolation would remove informative cues and thus harm individual recognition. The present study aims to adjudicate between these two possible scenarios.

## Materials & Methods

### Data

We studied five adult captive Japanese macaques (two males and three females) at the Primate Research Institute, Kyoto University (KUPRI), Japan. Before the experiment, the primates were kept in an outdoor holding cage in the same group of five). The macaques were then moved to an experimental area, which consisted of two outdoor cages of the same size (5 × 4 × 2.5 m; see Fig. 1). The two cages were adjacent to each other and partitioned by a wall with a gate. One of the cages was equipped to record the 3D location data of the macaques inside. We shut one or all the subjects in that cage by keeping the gate closed during the recording. Otherwise, if the gate was open, the subjects were allowed to freely move between the two cages. The macaques were fed primate pellets with supplemental vegetables once a day under the standard nutritional requirements for non-human primates.

**Figure 1:**
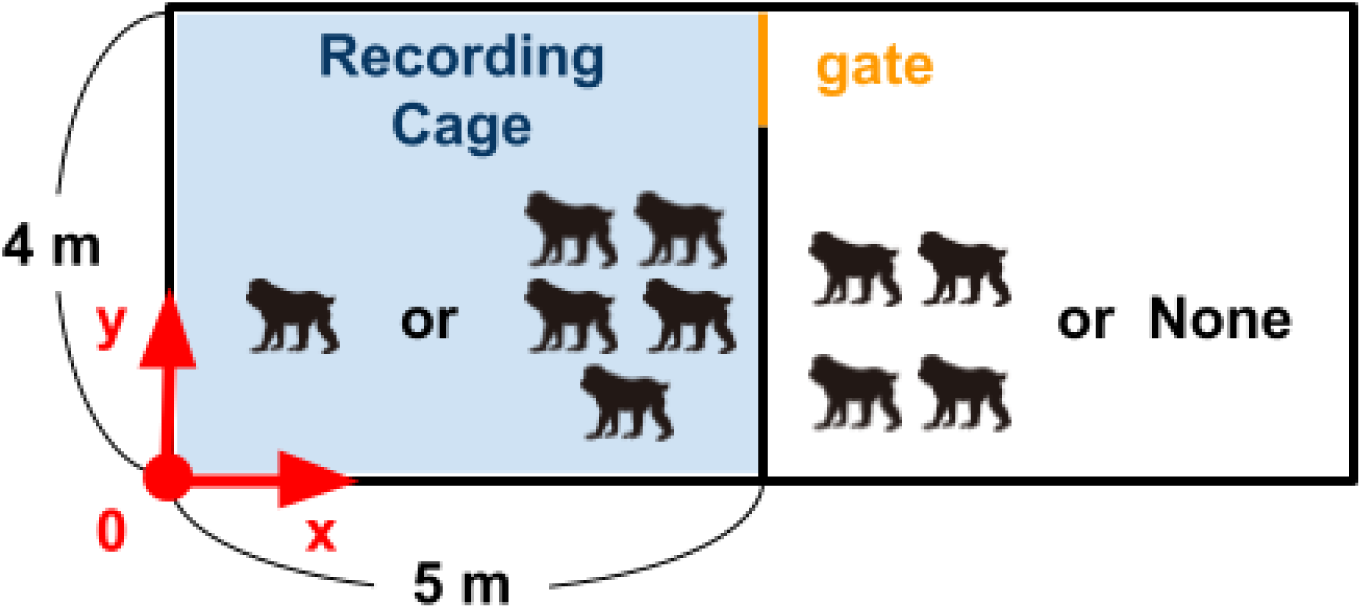
Recording environment. The 3D location data of one or all of the five macaques were recorded in the left cage; The other macaques were kept in the other cage on the right during the recording.

The 3D location trajectories of the macaques were recorded using a real-time tracking system based on Bluetooth^®^ Low Energy (BLE) beacons (Quuppa Intelligent Locating System™, AIBLE-QBP; 44 × 33 × 8 mm, 10 g). Each macaque carried three BLE beacons attached to a custom-made collar (approximately weighing 50–54 g in total). The signals from the BLE beacons were collected via six receivers. Four receivers were installed above the cage, and two on the *y* = 0 side of the cage (see Fig. 1). The system was designed to sample data in 5 Hz, but the actual rate in 1 second varied between 0 (no sample) and 6 Hz, peaking at 5 Hz (Fig. 2). Accordingly, we used the median coordinate values of the three beacons per individual over each 1-s window as an input data point for analyses (see the next section for more details).

**Figure 2:**
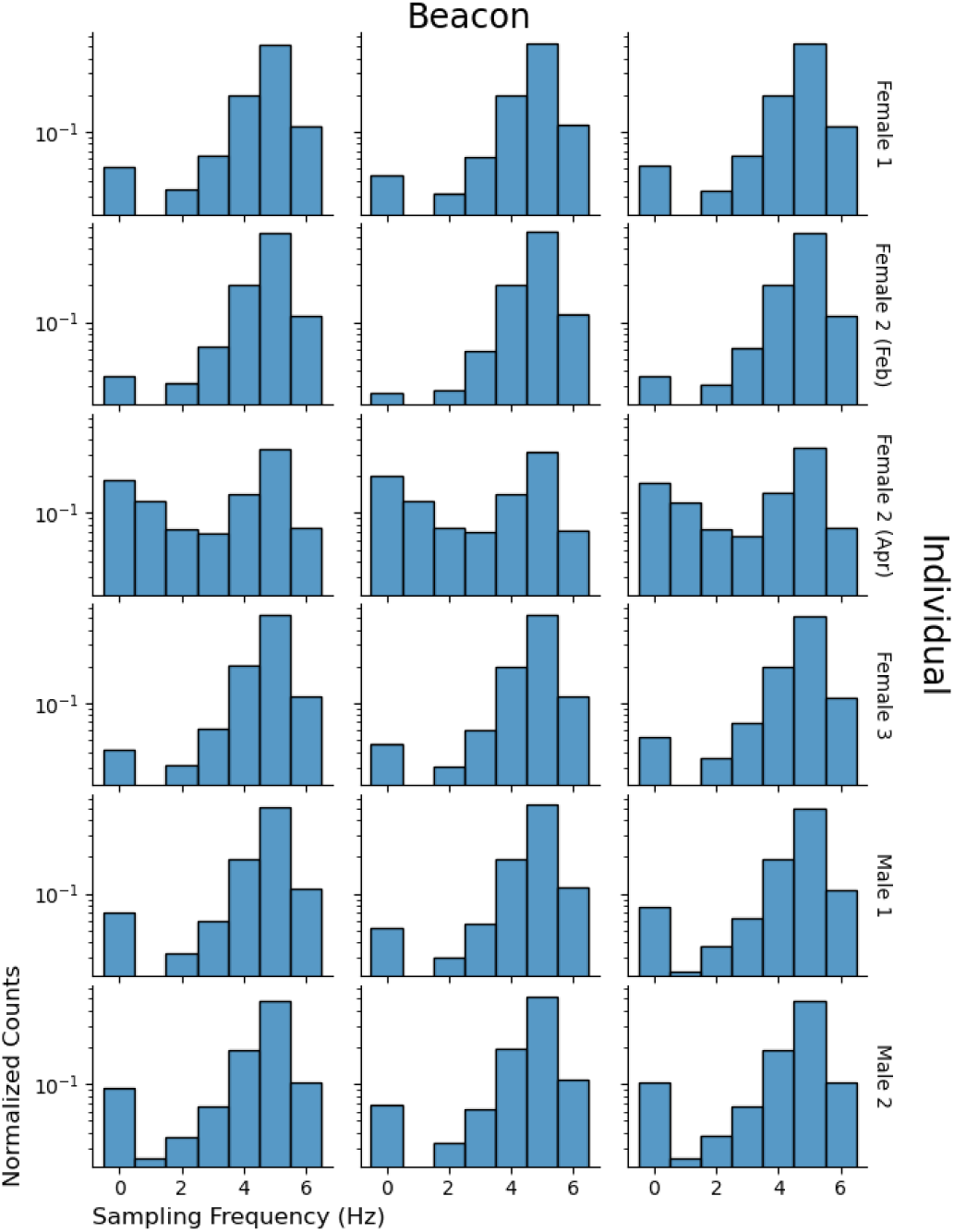
Empirical sampling frequency of Bluetooth^®^ Low Energy beacons. Each histogram reports the distribution of the sampling frequency from a single beacon. Three beacons in the same row were installed on the same individual macaque. Note that we replaced the beacons for Female 2 due to battery death, and, accordingly, data from two triplets (recordings from February and April) are reported for that individual.

Each of the five macaques was isolated for six to eight hours per day × four days (see Table 1 for the dates of the recordings). The isolation started between 9:00 and 10:00 by letting the non-target macaques move to the non-recording cage and then closing the gate between the two cages. The isolation lasted until the gate was reopened between 16:00 and 17:00. We analyzed the data between 10:00 and 16:00 on each day. The isolation was performed on weekdays, while all macaques were kept together in the recording cage during weekends, which allowed us to collect group-behaving data. Four non-rainy days were selected for the analysis from weekends between February 8 and 23, (most of the isolation data were collected on weekdays in the period). Only one of the twenty weekdays was rainy and thus we excluded the rainy weekend days from the analysis. Just like the isolation data, we used the 10:00-16:00 portion of the group-behaving data for the analysis. The battery of the BLE beacons died before we collected all the data as initially scheduled. This setback caused a delay of approximately one month between the last and penultimate days of the recording (April 7 for Female 2 and March 4 for Male 1, respectively) since it took about a month to replace them and start the recording again.

**Table 1:** Dates of the recording, all in 2020. For each day, we used the 3D location data of one (“isolated” condition) or all (“in-group” condition) of the five macaques that were collected between 10:00 and 16:00. The columns tell which one-hour portion of the day was used as the test data in the analysis.

| Subject Recorded | Condition | Portion of data used for the test |  |  |  |
| --- | --- | --- | --- | --- | --- |
|  |  | 11:00–12:00 | 12:00–13:00 | 13:00–14:00 | 14:00–15:00 |
| Female 1 | Isolated | Feb. 26 | Mar. 3 | Feb. 7 | Feb. 18 |
|  | In-Group | Feb. 15 | Feb. 23 | Feb. 8 | Feb. 9 |
| Female 2 | Isolated | Apr. 7 | Feb. 25 | Feb. 13 | Feb. 4 |
|  | In-Group | Feb. 23 | Feb. 15 | Feb. 9 | Feb. 8 |
| Female 3 | Isolated | Feb. 12 | Feb. 19 | Feb. 27 | Feb. 5 |
|  | In-Group | Feb. 9 | Feb. 15 | Feb. 23 | Feb. 8 |
| Male 1 | Isolated | Feb. 6 | Feb. 14 | Feb. 21 | Mar. 4 |
|  | In-Group | Feb. 8 | Feb. 9 | Feb. 15 | Feb. 23 |
| Male 2 | Isolated | Feb. 28 | Feb. 10 | Feb. 17 | Feb. 20 |
|  | In-Group | Feb. 23 | Feb. 8 | Feb. 9 | Feb. 15 |

### Analysis

Our primary question was whether the isolation caused any changes in the macaques’ 3D location trajectories compared to the in-group baseline. We addressed this question by computing the identifiability of the isolation vs. in-group condition from a location trajectory of a macaque under the condition. Specifically, we trained a convolutional neural network (CNN) [35–40] that took a five-minute location trajectory of a macaque as its input and made a binary prediction of whether the location trajectory was produced under the isolation or in-group condition (see Fig. 3). The input data were in 1 Hz, where each discrete time step represented the median of the location records collected from the three beacons in 1 second. The accuracy of the CNN’s predictions on the test portion of the data that were held out from the CNN’s training process defined the identifiability of the experiment conditions (specified in Table 1, amounting to 132,000 trajectories, 1/6 of the whole data). Hyperparameter values were borrowed from previous studies (see the supporting information S1 for details) and not systematically optimized on another held-out dataset for validation.

**Figure 3:**
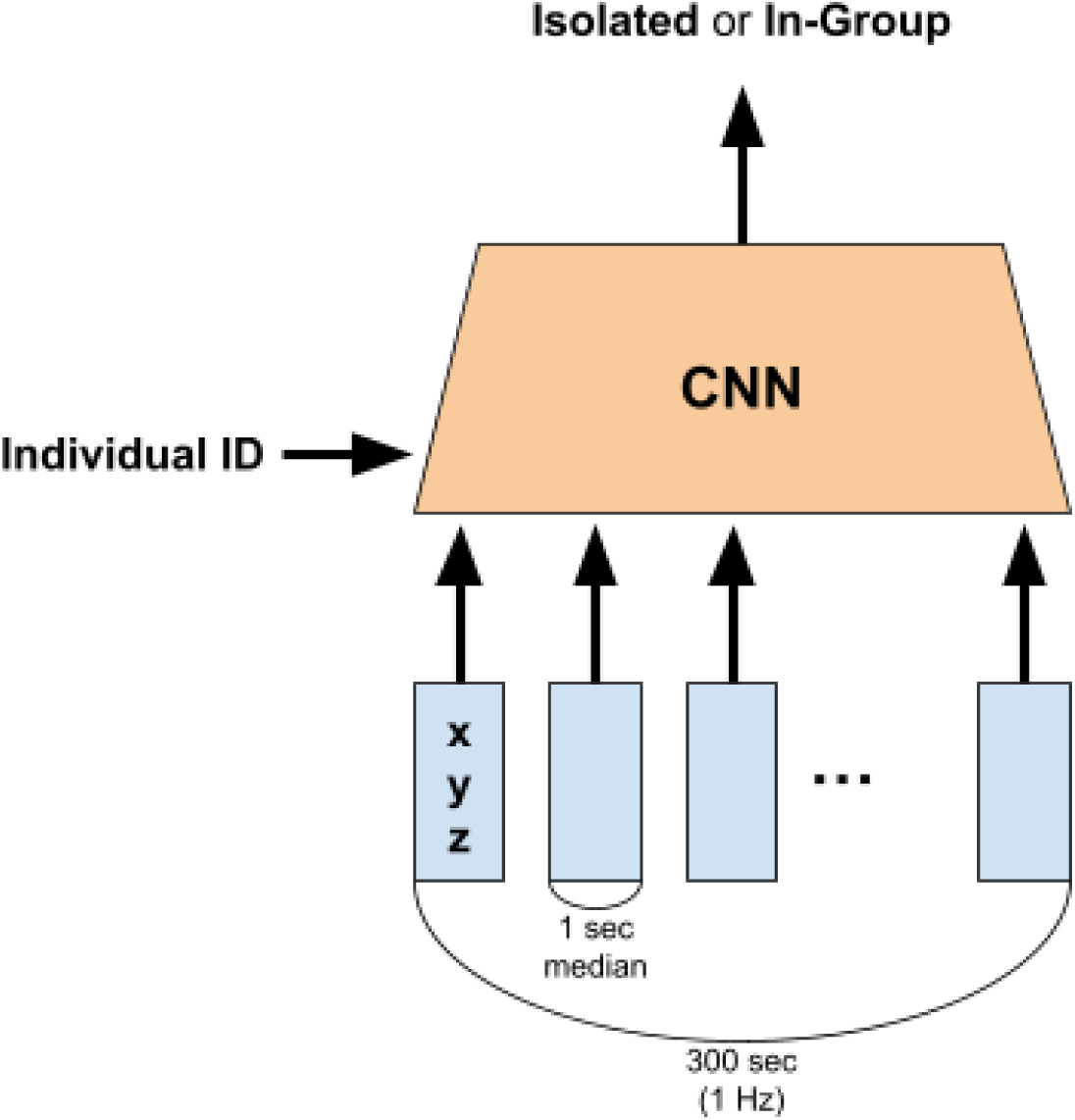
Overview of the model architecture. See the Supporting Information S1 for more detailed information about the CNN. The model described here predicted the experimental condition under which the input 3D location trajectory was produced. The model was also informed which individual produced the input trajectory. The individual identification was performed using the same model except that the output and background information were flipped (i.e., the individual identity was predicted from the location trajectory and the experimental condition).

To further diagnose the effect of isolation, we also assessed the identifiability of individual macaques from the location trajectories [25]. We used the same CNN architecture for this assessment, except that the network outputs the individual identity predicted as the agent that produced the input trajectory. We compared the prediction accuracy based on the isolation and in-group data.

It is to be noted that time-series data, such as human speech and text data, are often analyzed by a recurrent neural network (RNN) [41–43] whose input can have variable lengths in the time dimension (see also [44,45] for a more recent model of time-series processing and [25,46] for applications of RNNs to biological studies). In contrast, CNNs require their inputs to have the same length (when they return a single output [47–50] instead of converting the input to another time-series [51,52]). Despite this inflexibility, CNNs have an advantage over RNNs in greater interpretability of their inference thanks to the availability of effective visualization methods [32,53,54]. Inferences made by neural networks are, generally, difficult to interpret for human researchers because they result from compositions of many atomic numerical computations rather than a series of logical decisions. Thus, researchers need to make an extra effort to diagnose what kinds of information are used by the network, and there are visual explanation techniques exclusively available for CNNs’ inferences [32,53,54] besides model-agnostic methods [55–57]. In this study, we used Grad-CAM [32] (CNN-specific method) and SHAP [56] (model-agnostic method) to visualize specific segments of the location trajectories that were important for our CNNs to identify the experimental conditions and individual macaque behind the data.

The two visualization methods differ in their definition of importance score. Grad-CAM defines the importance of time steps based on the downsampled time-series data (1/12 Hz) through CNN. The downsampled signals, having 64 latent channels, yield the final classification probability directly by their weighted sum over time and channels; thus, they help determine which time step contributes to an increase in the probability. Since we are only interested in the importance of time steps, not of the latent channels, Grad-CAM computes the weighted average of the downsampled signals over channels, defining these “weights” by the gradient of the classification loss with respect to each time and channel of the downsampled signals. The resulting channel-averaged signals are clipped into non-negatives, upsampled into the original sampling rate at 1 Hz by linear interpolation, and then normalized into a 0–1 scale to yield the Grad-CAM importance score of each time step. Notably, Grad-CAM scores only show the importance of time steps within the input time series due to the final normalization step. Accordingly, we also weighted the Grad-CAM scores of each time step by the classification probability computed by the CNN to obtain a metric of importance for segments of different location trajectories. In contrast, SHAP can compute the importance of time steps in any layer of the network, and we applied it to the first linear transformation of the input time series before downsampling. Importing the concept of contribution from game theory, SHAP computes approximate Shapley values [58] of each element in the transformed input. Shapley values decompose the deviation of the model output value from the average output (corresponding to an increase in the probability of the predicted condition/individual) into a linear combination of the elements in the transformed input. We can interpret these coefficients, just as in linear regression, to find which elements increase the classification probability. Again, we were only interested in the importance of time steps, so we summed the SHAP values over the latent channels, computing the total contribution of each time step.

In addition to the main results based on CNN, we also report classification accuracy of three baseline models: logistic regression [59], support vector machine (SVM; with linear kernel) [60,61], and random forest [62] (see the supporting information S1.3 for their training procedures).

## Results

### Identifiability of Experimental Condition

Using the neural network-based classifier, the experimental condition―isolated or in-group―was identifiable from a five-minute location trajectory of the macaques with an accuracy of 92.71% (Table 2; 95% confidence interval, CI, ranged 92.57–92.85 according to 10,000 iterations of bootstrapping). This result suggests that isolation robustly changed the macaques’ behavioral patterns compared to the canonical group condition. Male 1 was the single subject that exhibited a notably lower accuracy (80.26%; 95% CI ranged 79.77–80.73) than the others (93.71%–98.75%), though this minimum score was still higher than expected by chance (=50%). This macaque was considered dominant over the other male (Male 2) according to a supplementary analysis of video recordings (24 h of observation, daytime on August 17–20, 2020); we observed five incidences of presenting behavior by Male 2 toward Male 1 and one incidence of mounting behavior by Male 1 over Male 2, which indicate the dominance of Male 1 over Male 2 [63].

**Table 2:**
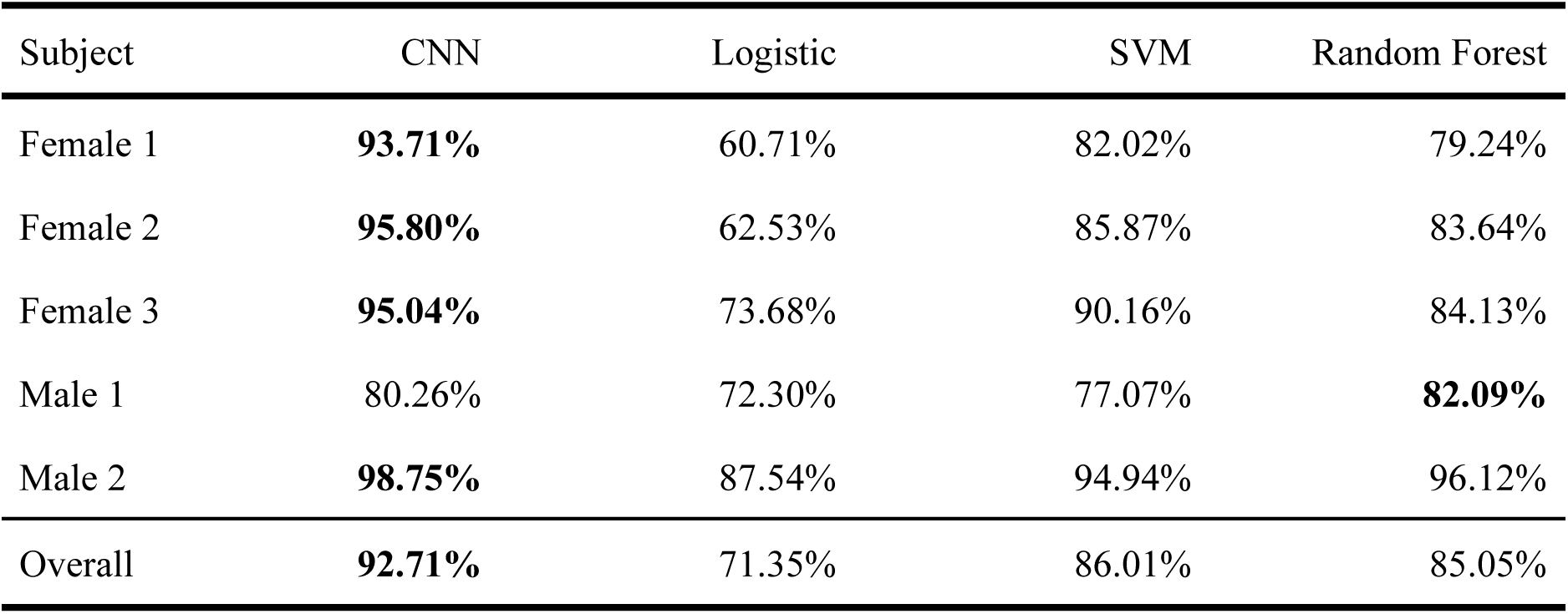
Accuracy of the condition classification based on the held-out test data.

| Subject | CNN | Logistic | SVM | Random Forest |
| --- | --- | --- | --- | --- |
| Female 1 | **93.71%** | 60.71% | 82.02% | 79.24% |
| Female 2 | **95.80%** | 62.53% | 85.87% | 83.64% |
| Female 3 | **95.04%** | 73.68% | 90.16% | 84.13% |
| Male 1 | 80.26% | 72.30% | 77.07% | **82.09%** |
| Male 2 | **98.75%** | 87.54% | 94.94% | 96.12% |
| Overall | **92.71%** | 71.35% | 86.01% | 85.05% |

The proposed classifier based on a neural network outperformed the baseline methods (Table 2; SVM was the best competitor for the overall classification; 95% CI ranged 85.82–86.19), except in the classification of Male 1 data (wherein the random forest achieved a slightly, but significantly better result; 95% CI ranged 81.63–82.55).

The identifiability evidenced robust differences between the macaques’ movements under isolated and in-group conditions. However, these were unrecognizable for human researchers (see Fig. 4A, B), until Grad-CAM visualization [32] of characteristic portions of the data helped locate where the differences reside (Fig. 4C). Specifically, it revealed that macaques in isolation were characterized by their staying around the wall that separates them from the other group members (*x* ≈ 5; see also Fig. 1 for the configuration of the experimental cage). This generalization was also supported by a different visualization based on SHAP (Fig. 4D).

**Figure 4:**
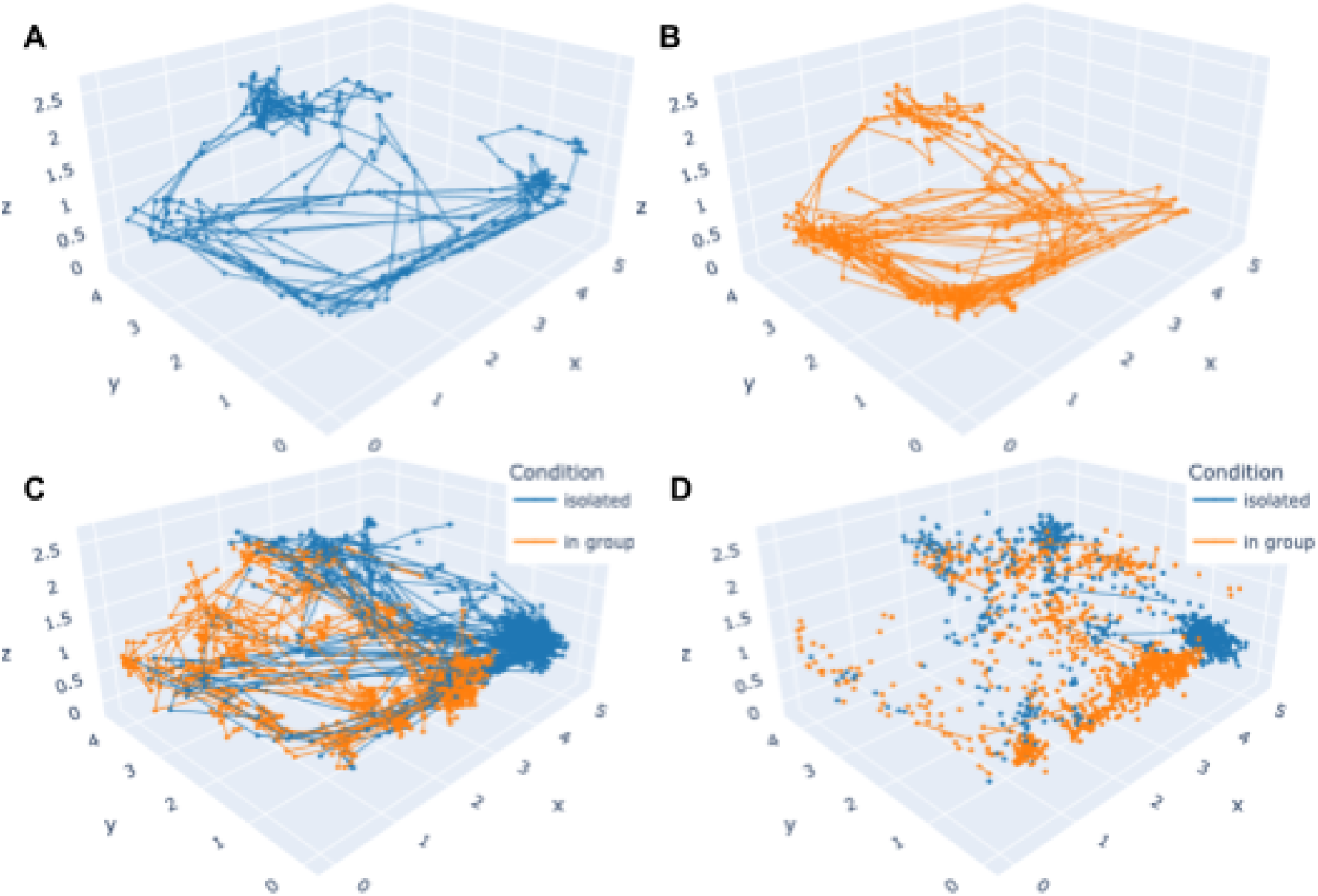
A five-minute location trajectory of a single subject (Male 1) under (A) isolation and (B) in-group conditions. The classifier identified the correct condition behind these trajectories with confidence, assigning ≥95% classification probability. (C) Grad-CAM visualization of the characteristic location trajectories of each condition [32], showing only the movement segments yielding ≥0.9 value of Grad-CAM score weighted by the classification probability. (D) SHAP visualization of the characteristic location trajectories of each condition [56] showing only the movement segments whose SHAP value is ≥99.9 percentile.

### Effects of Isolation on Identifiability of Individuals

The behavioral changes caused by the isolation also increased the identifiability of the individual subjects from their movements. Under the isolated condition, the CNN classifier recognized individual macaques in isolation with an accuracy of 87.44% (based on their five-minute location trajectories; 95% CI ranged 87.19–87.70), but, when they behaved in group, the accuracy was only 62.27% (Table 3; 95% CI ranged 61.90–62.65). This increase in individual identifiability was observed in all macaques and in most of the baseline results based on the other classifiers, though the accuracy was degraded overall in the baselines.

**Table 3:** Accuracy of the individual recognition based on the held-out test data.

| Subject | Condition | CNN | Logistic | SVM | Random Forest |
| --- | --- | --- | --- | --- | --- |
| Female 1 | Isolated | **87.44%** | 26.08% | 41.61% | 47.11% |
|  | In-Group | **45.89%** | 17.36% | 21.45% | 15.78% |
|  | Overall | **66.67%** | 21.72% | 31.53% | 31.44% |
| Female 2 | Isolated | **87.17%** | 15.23% | 58.30% | 48.76% |
|  | In-Group | **71.13%** | 14.11% | 26.81% | 32.05% |
|  | Overall | **79.15%** | 14.67% | 42.56% | 40.40% |
| Female 3 | Isolated | **92.92%** | 67.38% | 60.14% | 68.19% |
|  | In-Group | **72.94%** | 42.05% | 43.55% | 32.30% |
|  | Overall | **82.93%** | 54.71% | 51.84% | 50.25% |
| Male 1 | Isolated | **81.84%** | 20.59% | 23.78% | 20.75% |
|  | In-Group | 43.84% | 41.25% | 53.92% | 59.10% |
|  | Overall | **62.84%** | 30.92% | 38.85% | 39.92% |
| Male 2 | Isolated | **87.86%** | 6.95% | 81.44% | 80.13% |
|  | In-Group | **77.57%** | 11.91% | 51.22% | 66.92% |
|  | Overall | **82.71%** | 9.43% | 66.33% | 73.52% |
| Overall | Isolated | **87.44%** | 27.25% | 53.05% | 52.99% |
|  | In-Group | **62.27%** | 25.33% | 39.39% | 41.23% |
|  | Overall | **74.86%** | 26.29% | 46.22% | 47.11% |

Visualization of individual-characteristic movement did not yield an interpretable explanation (Fig. 5). Specifically, unlike characterization of the experimental conditions behind movements, Grad-CAM and SHAP showed inconsistent results, highlighting movements in different locations.

**Figure 5:**
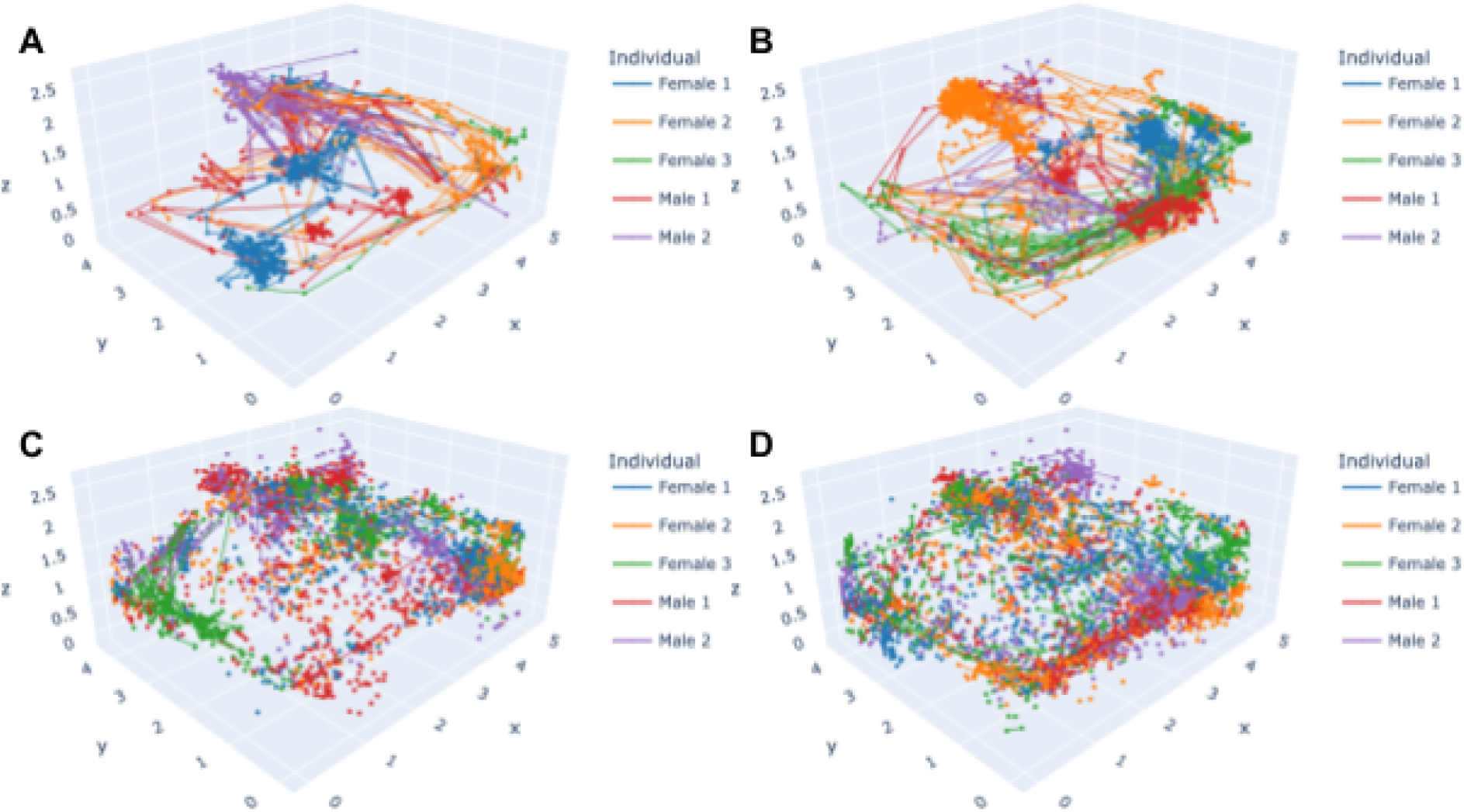
(A, B) Grad-CAM and (C, D) SHAP visualization of individual-characteristic location trajectories under the (A, C) isolated and (B, D) in-group conditions. Grad-CAM scores were weighted by the classification probability, and only the movement segments yielding a ≥0.9 weighted score are visualized. Likewise, only movement segments whose SHAP value is ≥99.9 percentile are visualized.

## Discussion

The present study compared location trajectories of captive Japanese macaques behaving in isolation vs. in group. The results demonstrated that the two experimental conditions were identifiable (with over 90% of accuracy) based on five-minute trajectories with the help of an artificial neural network. The result suggests that the macaques showed substantial changes in their movement patterns when isolated from other group members. Specifically, Grad-CAM visualization of characteristic movements for each condition revealed that the isolated macaques frequently visited the wall partition, where on the other side the other group members resided. A possible interpretation of this result is that the isolated individuals waited for the opportunity to rejoin the group. Our isolation kept the target individual in an experimental cage that neighbors another cage wherein the other non-target macaques were kept. Thus, although the isolated macaques could not see the others, they could still recognize the existence of the other macaques in the neighboring cage through non-visual information, such as sound and smell. For example, free-ranging Japanese macaques split up into small parties while keeping track of each other using their vocalizations [64]. Such vocal communication might have encouraged the isolated macaques to rejoin the main party in the next cage. Hence, future studies should consider such non-visual cues and regard more complete isolation beyond simple partitioning and further habituation to the different experimental conditions, while we acknowledge empirical difficulties with such refined experiments regarding ethical and management issues (e.g., using neighboring cages had the advantage of easy isolation and remerging). Despite the problem with experimental design, the discovered characteristic behavior under isolation was not easily identifiable in the raw location trajectories but was successfully visualized using machine learning techniques. Thus, the combination of experimental manipulation, large-scale data collection via biologging, and machine learning analysis is considered an effective research paradigm.

The notable exception was a male (Male 1) that showed a lower accuracy of the condition classification than other individuals, though the score was still far beyond the chance level (80.26% ≫ 50%). We suspect that this result is potentially related to the male’s social status in the group. Female-philopatric/male-dispersal species like Japanese macaques tend to exhibit greater spatial cohesion among females than among males [33,34]. Our previous study of the same group also showed greater statistical dependence among the females [26]. Thus, males are expected to move more freely from social constraints than females and be less influenced by the isolation, although the dominance hierarchy among males in the group still constrained their behaviors [65,66]. In addition, Male 1 was the dominant male and may have been allowed to move relatively freely even under the in-group condition, whereas the movement patterns of the other subordinates were more constrained in group. In short, thanks to his dominant social status, Male 1 need not change their movement patterns depending on the experimental conditions, which resulted in the lower accuracy of the condition classification.

The identifiability of individual subjects based on their movements increased when they were isolated. This tendency was observed across different classification models and, thus, it appears to be evident that some individuality traits emerged in location trajectory data under the isolation condition. Unfortunately, visualization of individual-characteristic movements did not yield an interpretable explanation about individuality, leaving issues to be addressed in future studies. A possible factor responsible for the difficulty in individual recognition under the in-group condition, from a behavioral ecology perspective, is the synchrony in activities of the macaques. Activity synchrony is one of the keys to enhancing group cohesion in group-living animals [67], and such synchrony has been widely reported in primates, typically in the context of feeding [68–70]. Behavioral synchrony upon feeding is expected to be more frequent in a captive environment than in wild situations. While preferred foods for free-ranging macaques (e.g., fruits and mushrooms) are seasonal and are distributed geographically patchily [71,72], captive feeding is regular. In this case, all the group members are fed approximately simultaneously, which provides a better opportunity for their behavioral synchrony. It is of note that behavioral synchrony can also occur in the form of social activities, such as grooming, but they are beyond the scope of the present analysis related to our location trajectory data. Thus, future studies are necessary to explore the loss of synchrony and other behavioral changes upon isolation or group split in other modalities, such as from video recordings.

Notably, the accuracy scores of individual recognition (based on the in-group data) were robustly greater than our previous study [25]. Several factors are considered responsible for this improvement. First, we used one additional beacon per individual in the current study. Thereby, the total sampling frequency per individual was higher than in the previous one, which only used two beacons per individual. This increased sampling frequency enabled us to use more stable data by taking the median of every second, while the previous study used the raw samples as the inputs to the model. Likewise, the median data contained a constant number of discrete time steps for each five-minute input, which allowed us to use the CNN-based classifier. The previous study, on the other hand, used an RNN-based classifier because the input time series contained a variable number of time steps. In this study, the model inputs were also longer in duration (5 min) than in the previous (2 min); therefore the model had a better chance of finding clues to identify individuals. Note that the data analyzed here were collected more than one year after the previous, and it is possible that there has been a change in the macaques’ behavioral patterns during the time.

From a technical perspective, it should be noted that neural networks are not the only option for analyzing large-scale biologging data. For example, recent studies propose graph-theoretic analyses of location trajectory data to discover interesting patterns in them [27–29]. Graphical analyses first quantize continuous-valued location data and associate the discretized locations with the graph nodes. Then, animal movements are represented by transitions between the nodes, and researchers can apply various analytical tools for graphs (e.g., centrality analysis) to the graphically represented movements. However, one major difference between neural network-based analyses and other methods, including graphical analyses, is the possible range of information aggregation over time. Taking the graphical analyses as an example, recall that a location trajectory is represented as a sequence of inter-node transitions. Then, statistical analysis of long-range movement patterns becomes hard due to data sparsity; the number of possible movement patterns grows exponentially as they get long, and almost all of them would become unique, having only a single observation. Accordingly, robust statistical inferences are limited to short-term windows consisting of only a few time steps [29]. This is similar to natural language processing (NLP) before the deep learning era; NLP was modeled by a Markovian process that can only integrate information from two to four words [73–75]. In contrast, neural networks are competent for integrating information from a long period (300 time steps in this study), exploiting real-valued latent feature spaces (even for discrete data, such as text languages) [76,77]. Thus, neural networks are particularly effective when researchers need an analysis of long-range time series or cannot deny their necessity. Moreover, neural networks have started to be applied to graphical data [78,79], which could potentially bring a novel insight into movement ecology as well.

As demonstrated here, the combination of behavioral experiments and modern technologies such as biologging systems (e.g., BLE beacons and GPS) and neural network analysis helps shed light on hitherto unrecognized latent factors/mechanisms behind behaviors of social animals. Future studies are necessary to explore a wider variety of split patterns (e.g., making subgroups of two or three members) to elucidate such complex social interactions.

## Acknowledgments

This study was funded mainly by the JST Core Research for Evolutional Science and Technology 17941861 (#JPMJCR17A4) and partially by MEXT Grant-in-aid for Scientific Research on Innovative Areas #4903 (Evolinguistic; JP17H06380). We appreciate the animal care given by our technicians and research assistants as well as their suggestions and support for this project. In particular, we wish to express special thanks to Norihiko Maeda, Mayumi Morimoto, and Takayoshi Natsume for the animal arrangements, and Panasonic Solution Technologies Co., Ltd. for the technical support with the equipment. This study was performed under the Cooperative Research Program at KUPRI (2018-C-27, 2019-B-27, 2020-B-17).

## Ethics Statement

All procedures were reviewed and approved by the Animal Welfare and Care Committee of KUPRI (Permission # 2018-203, 2019-208, and 2020-185) and complied with the institutional guidelines [80].

## Author Contributions

Project organization: IA IM HK; Animal arrangements: HK SA NSH; Apparatus building: AT IM HK; Data acquisition: AT IM HK; Animal cares: AK AT IM HK; Data management; HK AT TM; Computational modeling: TM; Manuscript writing: IA IM HK TM.

## Notes

### Competing Interest Statement

The authors have declared no competing interest.

