## Supporting Information for "Effects of short-term isolation on social animals’ behavior: an experimental case study of Japanese macaque"

#### S1 Detailed Information on the Proposed Method

##### S1.1 Neural Network Architecture

We built an artificial neural network which takes a 5-min location trajectory of a Japanese macaque as its input and returns a probability of the experimental condition under which the macaque exhibited the movement (either "isolated" or "in-group"). The network architecture is shown in Figure S1.1a. The network first applied a linear transform to each time step of the 3D location trajectory and represented them in a 512-dimensional feature space. The trajectory was in 1 Hz and the location coordinate $(x_t, y_t, z_t)$ at time $t = 1, \dots, T$ represented the median of the raw data collected from three beacons in the 1 s between $t$ and $t + 1$. Note that some of the 1 s time frames did not include any sample in the raw data since the empirical sampling rate of the beacons was not completely stable. Accordingly, the median vectors at the corresponding time steps had missing values, and we filled them with a trainable embedding in the 512-dimensional feature space after the linear transform (the same embedding was used for all missing values). We also embedded the ID number of the individual yielding the input trajectory in a 512-dimensional space, and additively combined it with the transformed trajectory. After activation by the leaky rectified linear unit (ReLU) and batch normalization (BN), the combined signal was downsampled from 1 Hz (in the input) to 1/12 Hz through three convolution blocks (Figure S1.1b). Each block first downsampled the signal by a strided convolution layer and, then, further transformed the downsampled signal with two non-strided convolution layers and two linear layers (each followed by the leaky ReLU activation, residual connection, and BN). The resulting 1/12 Hz signal was then reduced in the channel size (from 512 to 64) via a linear transform plus activation. Finally, a fully connected (FC) layer turned the signal into classification logits and the softmax operation normalized the logits into probability vectors over the two experimental conditions.

We used the same neural network for the individual recognition task except that the experimental condition was embedded instead of the individual's ID and the output probabilities were of the individuals rather than the conditions.

##### S1.2 Optimization

The neural network was trained for 8000 iterations to minimize the cross-entropy loss of the classification, using the Adam optimizer (Kingma and Ba, 2015). The batch size was 512. The beta parameters of the optimizer were set as $\beta_1 = 0.9$ and $\beta_2 = 0.999$, and the weight decay was 0.01. The learning rate was linearly warmed up from $0.001 \times 400^{-1.5}$ to $0.001 \times 400^{-0.5}$ in the first 400 iterations and, then, decreased according to $i^{-0.5}$ at the $i$-th iteration (Vaswani et al., 2017). The values on the hyperparameters were borrowed from previous studies (Devlin et al., 2018), except for the iterations (which were selected so that the training completed in a day; it took approximately 17.5 hours using a single NVIDIA RTX2080Ti card) and batch size (maximized under the memory constraint).

##### S1.3 Baseline models

In addition to the proposed method based on CNN, we also trained three baseline models: logistic regression (Cox, 1958), support vector machine (SVM; Vapnik and Lerner, 1963; Boser et al., 1992), and random forest (Breiman, 2001). Logistic regression was trained by stochastic gradient descent with the same batch size and iterations as CNN (512 and 8000 respectively). SVM and random forest were optimized by full-batch learning, using randomly selected subdata consisting of 10000 location trajectories. The maximum

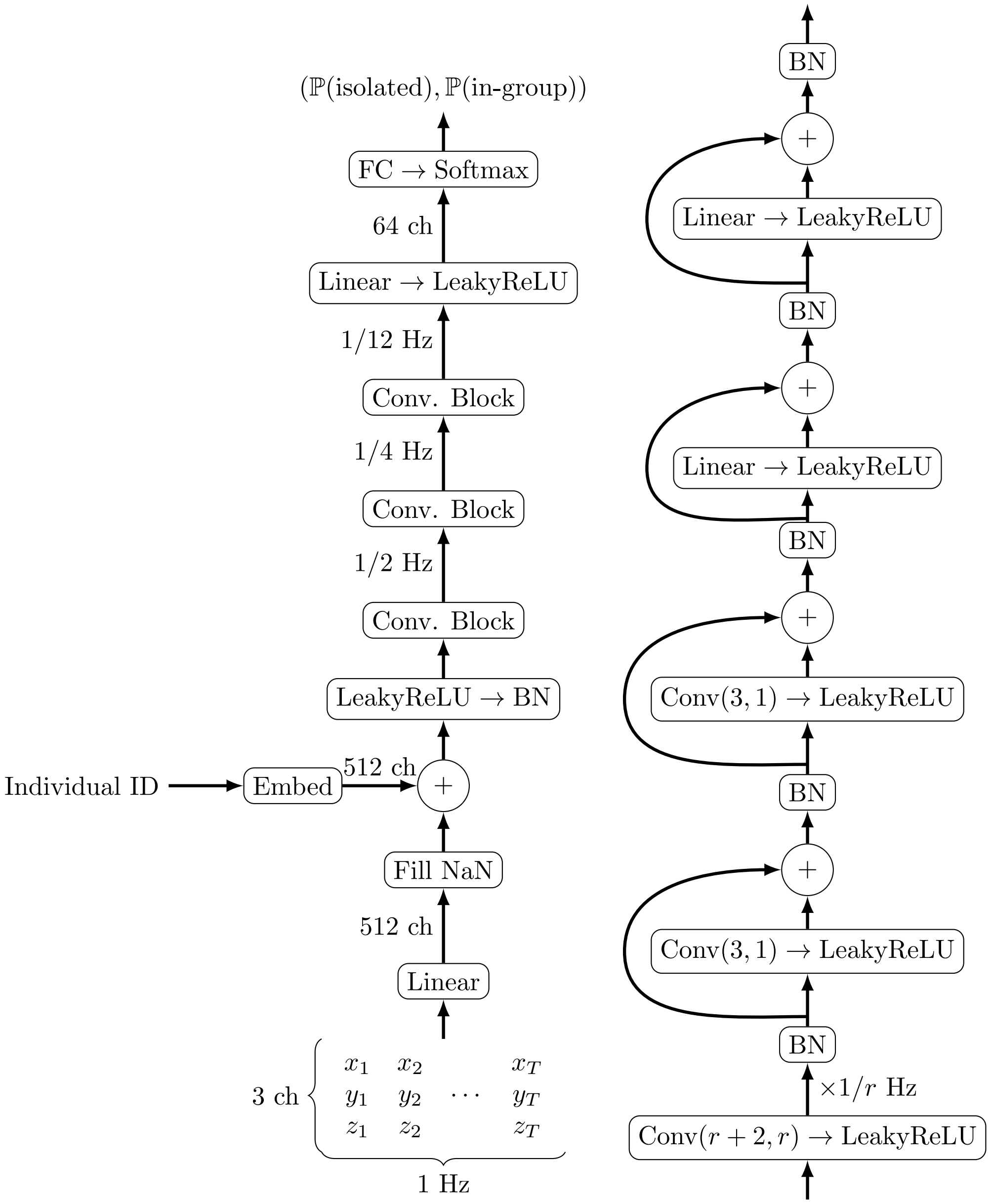


(a) Entire Network (b) Convolution Block

Figure S1.1: The architecture of the neural network adopted in this study. (a) The architecture of the entire network which computes the probability of the experimental conditions ("isolated" and "in-group") given a 5-min 3D location trajectory of a subject macaque and the individual ID of that subject. The network for individual recognition is obtained by swapping the individual ID in the (b) The convolutional block in which the time-series signals are downsampled by $r$ ($= 2, 2, 3$ respectively from the bottom) via convolution with a kernel of size $r + 2$ and stride of size $r$ (denoted by $\mathrm{Conv}(r + 2, r)$ in the figure; each convolution also zero-padded its input by one time step on both sides).

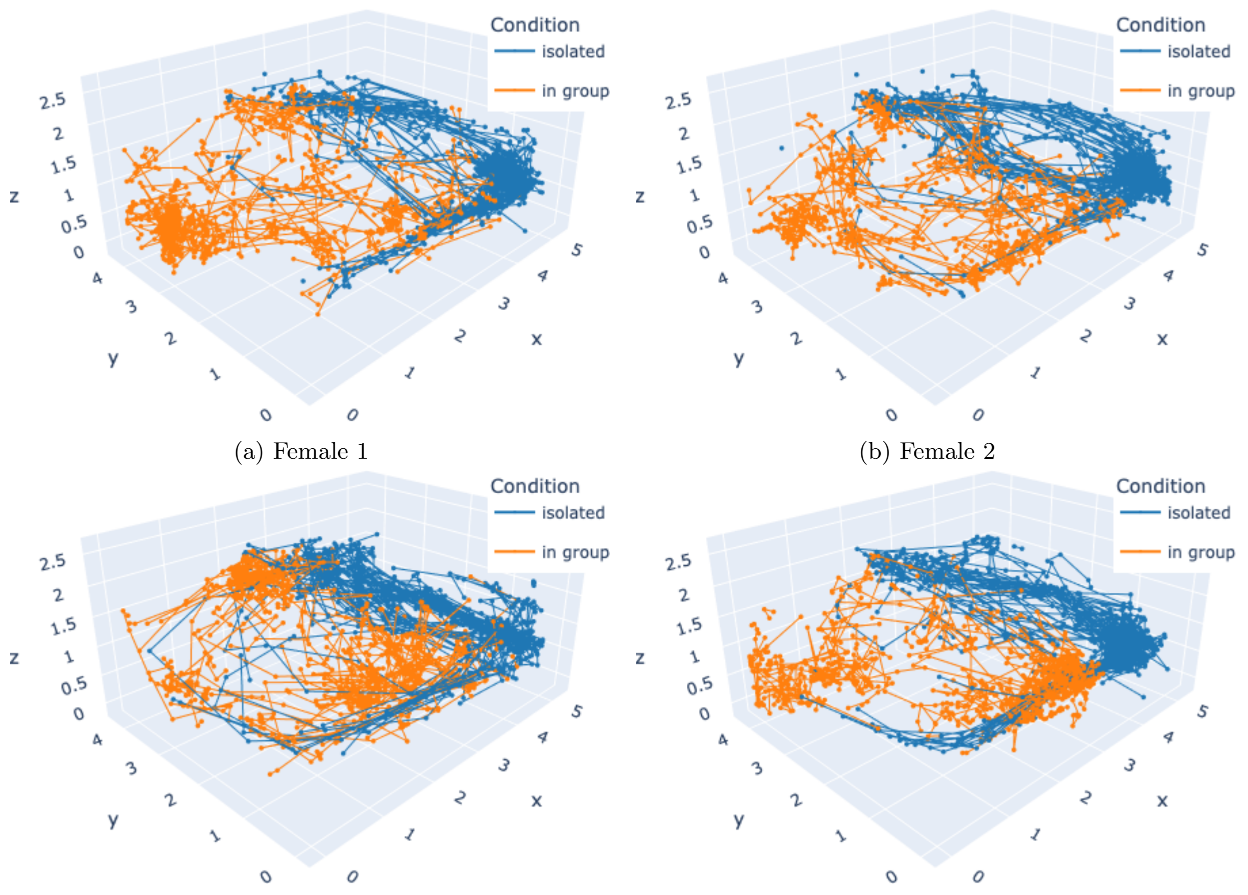


(a) Female 1
(b) Female 2
(c) Female 3
(d) Male 2

Figure S2.1: Grad-CAM visualization of the characteristic location trajectories of each condition (Selvaraju et al., 2017), showing only the data yielding 0.9 or a greater value of Grad-CAM score weighted by the classification probability.

iterations of SVM was set as 8000. The missing values in the data were filled with the nearest neighbor's value. We adopted scikit-learn's implementation (`sklearn.linear_model.SGDClassifier`, `sklearn.svm.SVC`, and `sklearn.ensemble.RandomForestClassifier`). The default hyperparameters were used unless otherwise noted.

### S2 Visualization of Condition-Characteristic Movements

Figure S2.1 reports Grad-CAM visualization of characteristic movements of the four macaques other than the one that is reported in the main text (i.e., Male 1; Selvaraju et al., 2017).
